# Seasonal and developmental stage changes in exudate carbohydrate content shape the kelp microbiome

**DOI:** 10.1101/2025.05.07.652744

**Authors:** Chance J. English, Meenakshi Manoj, Lillian C. Henderson, Keri Opalk, Craig A. Carlson

**Author notes:** Corresponding author: Chance J. English. **Author Contribution Statement**: CJE and CAC conceptualized the research. CJE collected material for and led the research. CJE, MM, LCH,and KO collected and analyzed data or performed laboratory measurements. CJE, MM, LCH,KO, and CAC contributed to interpreting the results and writing/editing the manuscript. CAC and CJE acquired funding for the research.

## Abstract

Photoautotrophs release a large amount of their fixed carbon as exudates, which shapes their immediate environment, including the composition of their microbiome. Here we evaluated the microbiome and exudate composition of *Macrocystis pyrifera* (giant kelp), a globally distributed foundation species, in response to seasonal nutrient availability and developmental stage. We combine 16S rRNA amplicon analysis of the giant kelp microbiome with carbohydrate monomer analysis of kelp exudates to examine microbe-exudate relationships. We found significant differences in the microbiome and exudate composition between seasons and developmental stages of giant kelp. Higher tissue-nitrogen content in the spring coincided with elevated amounts of glucosamine, a nitrogen-containing sugar, in giant kelp exudates, while senescence led to the shedding of mannuronic acid, an alginate indicator. The release of glucosamine and fucose-rich exudates was correlated with an increase in the relative abundance of bacteria within the Planctomycetota phylum, whereas mannuronic acid-rich exudates coincided with an increase in the relative abundance of members of the Flavobacteria and Gammaproteobacteria lineages. We investigated putative carbohydrate-microbe relationships by isolating a member of the Planctomycetota phylum from the surface of giant kelp. We demonstrate that the growth of this isolate on fucoidan and *N-*acetyl glucosamine, but not alginate reflects the observed relative abundance of this clade in the kelp microbiome in response to variable carbohydrate exudation. This suggests a key role of kelp mucilage carbohydrate composition in structuring its microbiome as has been observed for other organisms such as corals and within the human gut.

## Introduction

Primary producers such as plants and macroalgae have evolved in environments surrounded by microorganisms, and in many cases, associated microbes can improve the fitness and growth of the host through defense or nutrient acquisition [1–4]. A growing body of evidence indicates that autotrophic organisms regulate their microbiome by exuding metabolites that promote microorganism growth or inhibit the growth of pathogens [5, 6]. The exudate and host-microbe associations are dynamic and can vary in response to the host’s developmental stage and abiotic stressors [7, 8]. Unraveling the relationship between host physiology, exudate composition, and the microbiome structure can provide a mechanistic understanding of the establishment and function of the microbiome of primary producers.

Carbohydrates, including simple sugars, disaccharides and complex polysaccharides are typically the dominant organic material exuded by primary producers [9, 10]. The active exudation of carbon-rich organic matter by organisms was originally proposed as an “overflow” mechanism that allowed photosynthesis to continue when light and nutrients needed for biosynthesis became uncoupled [11, 12]. More recently, it has been proposed that the exudation of carbohydrate-rich material can also serve to prevent the attachment of pathogens [6], promote the growth of beneficial microbial partners [8, 13] or aid in the transport of energy and nutrients to the organism [14]. The potential role of carbohydrate exudation in shaping microbiomes is evidenced in studies of corals that reveal significant correlations between the host’s exuded mucus sugar content and the relative abundance of certain microbial taxa [15, 16]. The composition of exuded carbohydrates by corals is species-specific [17, 18] and varies in response to abiotic factors such as temperature [15]. While these relationships are established for corals, the role of carbohydrate exudation in shaping the microbiome has yet to be evaluated for important foundation species such as kelps.

Kelp forests are critical marine habitats that occupy 43% of the world’s coastline [19]. In addition to their contributions to biodiversity [20] and global economies [21], it is hypothesized that kelps contribute to carbon sequestration through the export of dissolved organic carbon (DOC) and detrital particulate carbon [22–24]. In this context, the decline of kelp forests [25] due to stressors such as ocean warming is a serious threat to ecosystem function, marine carbon cycling, and economies. This has prompted calls for large-scale intervention through natural kelp forest restoration [26] or aquaculture to offset these losses and increase current production yields [27]. Therefore, identifying and understanding the underlying factors for kelp health and resilience is of critical importance for monitoring, management, and restoration of these ecosystems, their services, and the economies dependent on kelp.

Decades of research have focused on understanding the impact of environmental factors such as light, nutrients, wave disturbance, and temperature on kelp growth [28–32]. In temperate coastal areas, such as California, kelp growth rates and physiology are influenced by seasonal nitrate availability. Low seawater nitrate concentrations (NO_3_^−1^ < 1µM) throughout the summer and fall result in elevated tissue carbon to nitrogen ratios and when tissue nitrogen content falls below 1% of dry mass, kelp survival can become compromised [32]. In comparison to abiotic factors, less is known about how the kelp microbiome affects kelp growth and resilience, although there is a growing appreciation of role of the microbiome in kelp health and resistance to disease [33, 34]. Several studies demonstrate that kelp harbor a microbiome that is distinct from the surrounding seawater [35–37] and meta- and whole genome analysis of kelp microbiomes suggest potential positive interactions, such as antibacterial and antiherbivore defense, or vitamin acquisition [38–40], between kelp and specific microbial lineages such as Planctomycetota. Further, there are significant associations between environmental conditions (i.e. temperature, nutrients), the physiological condition of kelp, its productivity, and its microbiome composition [41–43]. However, the underlying mechanisms between kelp physiology and microbiome composition are unresolved, although previous studies suggest exudate composition, environmental perturbations, or the interaction between the two may be important [44–46].

In this study, we aimed to understand how the composition of kelp exudates shapes the microbiome of *Macrocystis pyrifera*, hereafter giant kelp. We hypothesized that abiotic factors (nitrogen availability) and developmental stage would alter the chemical composition of kelp exudates and that these altered chemical profiles would shape the giant kelp microbiome. We sampled tagged cohorts of giant kelp fronds to track the development of their microbiome across developmental stages and between the warm, nitrate depleted summer and nitrate replete spring upwelling periods. Simultaneous measurements of exudate carbohydrate composition and 16S rRNA gene analysis were employed to identify microbe-exudate associations. We observed significant correlations between abundant microbial taxa and the carbohydrate content of giant kelp exudates. To test some of these associations, we isolated a member of the Planctomycetota phylum, a dominant kelp-associated bacterial group. Using whole genome sequencing, phylogenetic analysis and growth assays, we demonstrate that the carbohydrate substrate preferences of this clade reflect its relative abundance patterns in the kelp microbiome. This work suggests an important role for mucilage carbohydrate composition in shaping the kelp microbiome.

## Methods

### Sampling, kelp blade age and developmental stage, and environmental variables

Giant kelp blades were sampled from two tagged cohorts at Mohawk Reef (34° 23.6600’ N, 119° 43.8000’ W) in the Santa Barbara Channel, CA, USA in the summer of 2023 and spring of 2024. Cohorts were tagged by placing a cable tie around the stipe 2 m back from the apical meristem of 100-200 growing fronds. At intervals of 2-3 weeks, single blades (6 – 32 g wet weight) from 6 random fronds were sampled to measure their tissue nitrogen (tissue-N), exudate carbohydrate composition and microbiome. The developmental stage of kelp blades was categorized as either “mature” or “senescent” if they were younger than or older than 50 days old, respectively [29]. Blades were collected in the morning (08:00-10:00) and transported in a plastic cooler filled with ambient seawater to a near-shore laboratory, where they were incubated in 10 L acrylic tanks filled with filtered (0.2 µm) seawater. Incubations were started between 10:00-12:00 and maintained at *in situ* temperature (Summer: 17-19 °C, Spring: 12-14 °C) using a temperature-controlled water bath for 6-9 hours. Incubation tanks were equipped with magnetic stir bars to maintain water circulation. Light levels were adjusted every 2-3 hours, decreasing from saturating intensity (1140-1517 µmol photons m^−2^ s^−1^) to darkness (0 µmol photons m^−2^ s^−1^) to simulate the transition from day to night. Duplicate DOC (n = 84) and one hydrolyzable carbohydrate (n = 42) sample were collected by siphon at the beginning and end of each incubation and filtered through pre-combusted glass-fiber filters (Whatman GF-75, 0.3 µm nominal pore size). Following their incubation, kelp blade tissue-N was sampled as a measure of nitrogen stress and developmental stage. For tissue-N analysis, giant kelp blades were scraped cleaned of epibionts with a metal spatula, dried at 60 °C for three days and then ground to a fine powder before analysis of a 10 mg subsample with an Exeter Analytical CE-440 CHN/O/S elemental analyzer.

Environmental conditions at Mohawk Reef during our study were evaluated using bottom temperature data collected by the Santa Barbara Channel Long-Term Ecological Research program [47]. Average daily temperature was calculated from submersible temperature loggers at Mohawk Reef. Nitrate concentrations were calculated from temperature using an exponential “T2N” relationship derived from an inshore temperature and nitrate dataset [48] (Supplemental Figure. 1).

**Figure 1.**
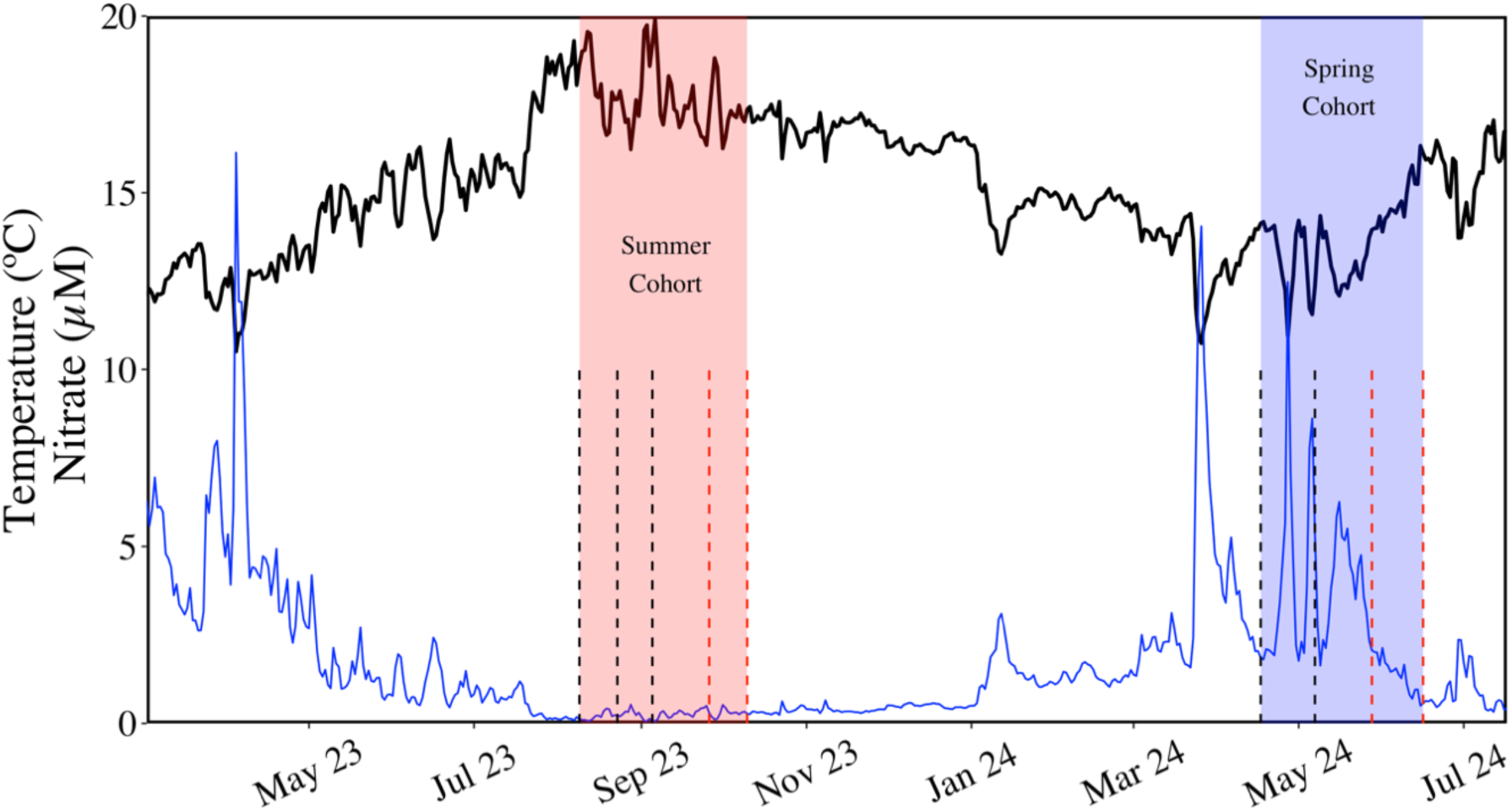
Mohawk Reef environmental conditions and sampling dates in this study. Daily average temperature (solid black line) and nitrate (solid blue line) concentrations between January 2023 and August 2024. Shaded red and blue regions show the sampling periods for the summer and spring cohorts used in this study, respectively. Vertical dashed lines extending from the x-axis mark the sampling dates within each cohort. The black and red color of the dashed lines indicate sampling of kelp that was in a mature (< 50 days) or senescent (> 50 days) development stage, respectively.

### Exudate DOC, carbohydrate content and composition

Dissolved organic carbon (DOC) analysis was carried out following established protocols [49]. Briefly, duplicate DOC samples were taken from at beginning and end of each incubation, filtered through pre-combusted 25 mm GF-75 into pre-combusted 40 ml EPA vials with PTFE lined caps, acidified to pH < 3 with the addition of 60 µl of 4N HCl and analyzed using a Shimadzu TOC-V or TOC-L.

Dissolved carbohydrate samples were taken alongside DOC samples and stored frozen at -20°C. Exudate carbohydrate content was measured by high-performance anion exchange chromatography with pulsed amperometric detection (HPAEC-PAD). We followed established dialysis and HPAEC-PAD gradients [50] . Briefly, samples were dialyzed using Spectra/Por 7 tubing (1000 Da) against ultrapure water and then hydrolyzed in duplicate for 20 hours at 100°C in 0.4 M HCl then neutralized under a flow of N_2_ gas. Samples were run on a DIONEX ICS5000+ and sugars were separated using a Carbopac PA10 column (4×250mm) with a Carbopac PA10 guard column (4×50mm). Neutral and amino sugars were eluted with 18mM NaOH followed by 100mM NaOH/200mM Na-Acetate to elute acidic sugars. The system was calibrated using a common standard sugar mix [17, 51] dissolved in ultrapure water containing fucose (1mM), rhamnose (1mM), arabinose (1mM), galactosamine (0.25 mM), glucosamine (0.25 mM), galactose (1mM), glucose (1mM), mannose+xylose (2 mM), galacturonic acid (1 mM), glucuronic acid (1 mM) and mannuronic acid (0.5 mM). Linearity of the calibration curves were observed for concentrations ranging from 10nM-1µM. In our dissolved carbohydrate samples, we observed glucose and mannose+xylose contamination due to leaching from the Spectra/Por 7 dialysis tubing, so these sugars were removed from subsequent analysis. Previous studies demonstrate these sugars are not substantial components of carbohydrates released by brown macroalgae [17, 52].

Principal component analysis was used to visualize changes in the composition of hydrolyzable sugar monomers in giant kelp exudates. Sugars were normalized to molar percentages (mole%) relative to other sugars. Statistical analysis of the differences in the composition of the carbohydrates released by giant kelp between the two seasons and developmental stages was evaluated by PERMANOVA using the *adonis* function in R (vegan 2.7-1) [53] on the scaled mole% of individual sugar monomers.

### Microbiome DNA extraction, 16S rRNA gene sequencing & amplicon analysis

Following their incubation, but before their preparation for tissue-nitrogen analysis, kelp blades were rinsed with 0.1 µm filtered sterilized seawater and sampled across the entire surface by scraping with sterile cotton-tip swabs (Puritan®) for 30 seconds. Swabs were placed in sterile 2 ml cryogenic tubes and frozen at -80 °C. DNA samples were lysed in sucrose lysis buffer (750 mM Sucrose, 20 mM EDTA, 400 mM NaCl, 50 mM Tris-HCl, pH = 8.0) with 1% SDS and 0.2 mg mL^−1^ proteinase-K at 37°C for 30 minutes for cell lysis and protein digestion, then at 55°C for 30 minutes to complete the lysis and degradation of cellular components. DNA was purified using the phenol-chloroform method [54]. For amplification, we targeted the V4 region of the 16S rRNA gene using the 515F-Y (5’-GTGYCAGCMGCCGCGGTAA-3’) and 806RB (5’-GGACTACNVGGGTWTCTAAT-3’) primers (predicted 84% coverage of 16S rRNA sequences; SILVA TestPrime 1.0) with custom adaptors [55]. PCR was carried out in 25 µL reactions with 12.5 µL KAPA Robust Hotstart ReadyMix (final concentrations: 0.2 mM each dNTP, 1U DNA Polymerase, 2mM MgCl_2_), 1uL each of the forward and reverse primers (final concentration: 0.4 µM each), 2µL of bovine serum albumin (final concentrationl: 1.6 µg µL^−1^), 6.5 µL of PCR-grade water and 2µL of sample DNA and cycled for 3 minutes at 95 °C, followed by 30 cycles of 30 seconds at 95 °C, 30 seconds at 57 °C and one minute at 72 °C, ending with 10 minutes at 72 °C. Four negative controls (PCR-grade water) and two mock community samples were included in the PCR run for quality control. Following PCR, amplified DNA was quantified (Qubit, Thermo Fisher) and manually normalized by pooling equal nmol amounts of DNA from each PCR reaction well. Pooled DNA was concentrated using Amicon® Ultra 0.5mL 30K centrifugal filters (Millipore) then non-target DNA bands were removed by gel electrophoresis and extraction. Samples were sequenced on an Illumina MiSeq PE250 and demultiplexed, with zero barcode/index mismatches allowed, at the University of California, Davis DNA Technologies Core.

Demultiplexed raw sequence reads were quality checked and processed using DADA2 [56] in R. Forward and reverse reads were trimmed to 240 and 150 bp, respectively. Trimmed and error corrected reads were dereplicated and pair-end reads were merged and assigned taxonomy using the SILVA database [49; v138.1 with species]. In all samples, amplicons identified as mitochondria (0-100 reads, mean = 18 reads) or chloroplasts (143-20,725 reads, mean = 3,216 reads) were removed. PCR-water and mock community controls were checked to confirm consistent amplification and lack of contamination and then removed from further analysis (Supplemental Data 1). Quality filtering of chimeric sequences or sequences with ambiguous bases resulted in 4.4×10^6^ sequences from 54 samples. Samples had a read depth from 25,077 to 106,379 sequences, averaging 81,976 and were not rarefied for further analysis [58]. We analyzed differences in microbially community composition on kelp blades by season (summer vs. spring) and developmental stage (mature vs. senescent).

For multivariate analysis, singleton and doubleton amplicon reads were removed and remaining amplicon sequence variants (ASVs) were converted to relative abundances and normalized using a variance stabilizing transformation. For beta diversity calculations, samples were assigned NMDS scores based on Bray-Curtis dissimilarities using the *metaMDS* function in phyloseq [59] and differences in community composition between seasons and developmental stages were tested using a PERMANOVA with the *adonis* function in the R [53] . For Shannon diversity calculations, performed with phyloseq, singleton and doubleton amplicons were not removed. Significant differences in the mean relative abundances of amplicons by season and developmental stage were evaluated using the “DEseq2” package in R [60]. Differentially abundant ASVs were those that displayed a log2 fold-change >1.5 between developmental stage or season and a significant false discovery rate (FDR)-adjusted p-value (<= 0.01).

To identify relationships between the giant kelp microbiome and the exudate carbohydrate composition we used hierarchical clustering of Spearman’s rank correlations between the relative abundance of significantly differentially abundant ASVs and the mole% of sugar monomers simultaneously released by giant kelp blades. This clusters ASVs based on their degree of correlation with the sugar content of kelp exudates and allowed us to identify positive or negative microbe-carbohydrate relationships. Hierarchical clustering was arranged using Ward’s minimum variance method (squared Euclidean distance) with the *pheatmaps* [61] package in R.

### Planctomycetota cultivation, genome sequencing & growth on model carbohydrates

We enriched and isolated a strain of Planctomycetota following the suggestions of Lage & Bondoso (2012) and Wiegand et al., (2020) [62, 63]. Agar plates were prepared with 0.1 µm filtered seawater with 1% agar. Seawater was autoclaved with agar then supplemented with 1 ml L^−1^ vitamin solution (Biotin: 2 mg L^−1^, Folic acid: 2 mg L^−1^, Pyridoxamine-HCl: 10 mg L^−1^, Thiamine-HCl x 2H_2_O: 5 mg L^−1^, Riboflavin: 5 mg L^−1^, Nicotinic acid: 5mg L^−1^, D-ca-pantothenate: 5 mg L^−1^, Cyanocobalamine: 0.1 mg L^−1^, p-Aminobenzoic acid: 5 mg L^−1^, Lipoic acid: 5mg L^−1^, KH_2_PO_4_: 900 mg L^−1^) and 1ml L^−1^ trace metal solution (H_3_BO_3_: 2860 mg L^−1^, MnCl_2_ x 4H_2_O: 1810 mg L^−1^, FeCl_3_ x 6H_2_O: 316 mg L^−1^, ZnSO_4_ x 7H_2_O: 222 mg L^−1^, Na_2_MoO_4_ x 2H_2_O: 390 mg L^−1^, CuSO_4_ x 5 H_2_O: 79 mg L^−1^, Co(NO_3_)_2_ x 6H_2_O: 49 mg L^−1^). For a carbon and nitrogen source we added 10 ml of 5% *N*-acetyl glucosamine. To prevent fungal growth and select for Planctomycetota, the medium was supplemented with 20 mg L^−1^ of econazole nitrate, 200 mg L^−1^ampicillin and 1 g L^−1^ streptomycin. Bacterial inoculum was scraped from the surface of a giant kelp blade using a sterile swab for 30 seconds and resuspended in sterile medium without agar. The cell suspension was filtered through a 5.0 µm filter and enumerated using a GUAVA easycyte HT flow cytometer (Millipore). To each agar plate we added ∼2000 cells. Cell suspensions were streaked across the agar plates and incubated at 20 °C until colonies were formed. After 1 month, colonies showing characteristics of Planctomycetota (pink pigmentation, budding cell growth) were selected for restreaking. Candidate colonies were restreaked three times to ensure isolation and then identified using 16S rRNA gene metabarcoding of the V4 region using the SILVA Ref NR SSU r138.1 database. Whole-genome sequencing was performed on the Illumina NovaSeq X Plus platform, producing 2 x 151 bp paired-end reads (Seqcenter, Pittsburgh, PA). Trimmomatic (v0.39) and RQCFilter (BBTools v38.22) were used to remove low-quality or short reads. Reads were assembled and annotated with SPAdes (v3.15.3) and Prokka (v1.14.5), respectively, in the KBase platform [64]. Completion (99.93%) and contamination (1.18%) was checked using CheckM v1.0.18. For phylogenetic analysis, 14 publicly available isolate genomes from the *Pirellulaceae* family were selected from NCBI’s GenBank database and compared to our assembled genome (Supplemental Data 2). The genome of *Planctomicrobium piriforme* from the Planctomycetota order Planctomycetales was selected as the outgroup. A maximum-likelihood tree was constructed with GToTree [65] using 74 bacteria marker genes and visualized using the iTOL server. Carbohydrate-active enzymes associated with fucoidan and alginate degradation were predicted for our isolate and reference isolates using a set of Hidden Markov Models (HMMs) from the dbCAN2 CAZy collection using HMMER 3 software [66, 67]. Secondary metabolite production potential was analyzed using AntiSMASH [68].

We tested the ability of the *Rhodopirellula sp*. isolate to grow on model carbohydrates in batch cultures. We prepared artificial seawater (ASW) as incubation medium (0.42M NaCl, 0.029M Na_2_SO_4_, 0.01M KCl, 1mM KBr, 0.05mM H_3_BO_3_, 0.07mM NaF,0.05M MgCl_2_, 0.01M CaCl_2_, 0.06mM SrCl_2_, 2.5mM NaHCO_3_). Medium salinity and pH were adjusted to 33 ppt and 7.5, respectively and then autoclaved. Sterile ASW was amended with 0.1 µm filter sterilized vitamin and trace metal solutions (described above, 1ml L^−1^) and inorganic nutrients (10 mg L^−1^ NH_4_Cl, 1 mg L^−1^ KH_2_PO_4_). Triplicate 50 ml incubations were prepared for the following treatments: no amendment, *N*-acetyl glucosamine (NAGA; 25 mg L^−1^ ), alginate (25 mg L^−1^), and fucoidan (25 mg L^−1^). Because commercially available fucoidan is typically contaminated, notably by alginate and polyphenols [69, 70], we used ion exchange chromatography (IEX) to purify fucoidan sourced from giant kelp (Sigma-Aldrich, product #F8065). Purification was performed using a medium-scale IEX setup described by Sichert et al. (2021) [70]. The composition of the purified fucoidan was checked by ^1^H-^13^C HSQC NMR and HPAEC-PAD (Supplemental Figures 2, 3 & 4). Alginate (Sigma-Aldrich PHR1471) and NAGA (Fisher Scientific ICN100068) were certified to be 88 and 100% pure, respectively. An isolate colony was resuspended in sterile ASW without amended carbohydrates. After gently shaking for 20 minutes, the cell suspension was 5.0 µm filtered and enumerated using a GUAVA easycyte HT flow cytometer (Millipore). Cells were added to incubation vessels at a concentration of ∼6.0×10^4^ cells ml^−1^. For all treatments, triplicate control incubations without cells were sampled to confirm sterility of the incubation media and carbohydrate amendments. Incubations were monitored semi-daily by cell enumeration for nine days by flow cytometry (GUAVA easycyte HT, Millipore).

## Results

### Kelp physiological response to seasonal nitrate availability and developmental stage

We observed variations in kelp developmental stage driven by seasonal nitrate availability and age. In both seasons, rapid changes in physiology occurred after kelp blades passed 50 days of age, which we use to demarcate the transition from maturity to senescence. Across all samples, tissue-N (g_Nitrogen_/g_Dry Weight_ *100) ranged from 0.63% to 3.7%. Mature kelp sampled in the spring had a significantly higher tissue-N content than mature summer kelp due to higher seawater nitrate concentrations (Figure 1, Figure 2A; Welch’s t-test: t = -22.6, df = 19.0, p < 0.001). The average (±1 SD) tissue-N of mature kelp blades in the summer and spring were 0.92 ± 0.08% and 2.6 ± 0.14%, respectively (Figure 2A). After entering the senescent phase, average (±1 SD) tissue-N decreased to 0.7 ± 0.06% and 2.1 ± 0.77% in the summer and spring, respectively (Figure 2A).

**Figure 2.**
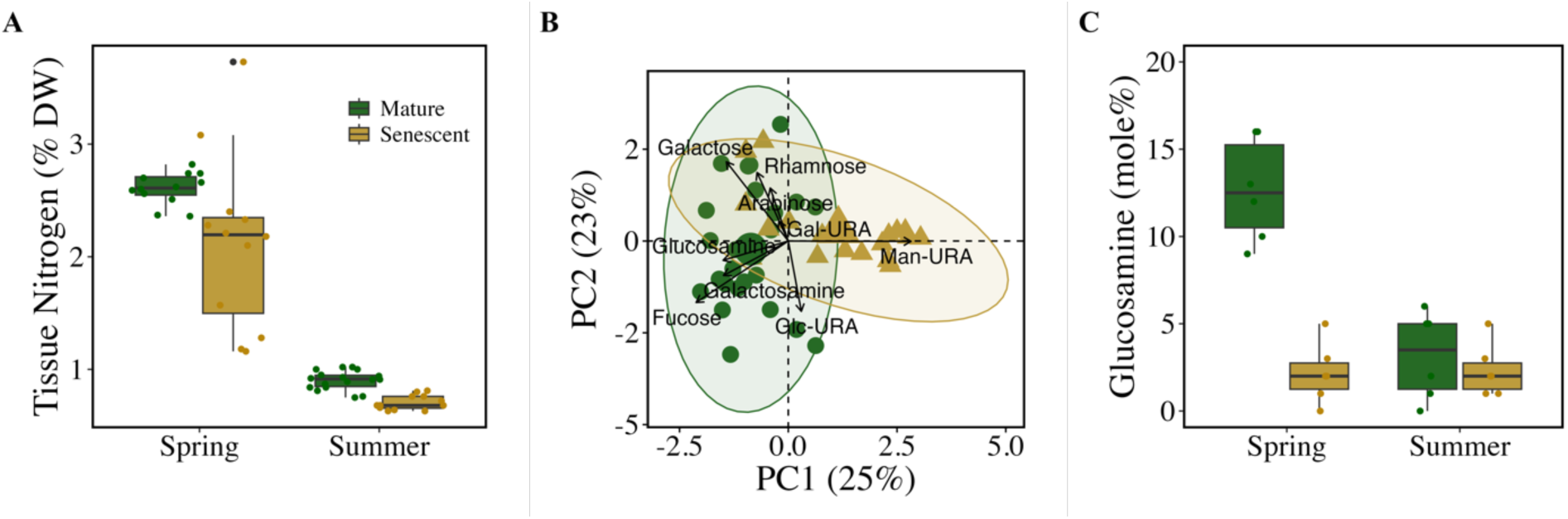
Seasonal and developmental stage changes in giant kelp physiology and exudate carbohydrate composition. **(A)** Giant kelp tissue nitrogen content as a percentage of dry weight (DW) between mature and senescent phase kelp in spring and summer. **(B)** Principle component (PC) analysis of giant kelp carbohydrate exudates as molar percentages between mature and senescent phase kelp. Ellipses represent 95% confidence regions between mature (green circles) and senescent (gold triangles) kelp exudates. Arrow lengths represent the strength of the correlation between each individual sugar monomer to the two principal components (PC1 & PC2) shown. Large symbols in center of the ellipses are the centroids. Sugar monomer names are overlayed next to arrows. Abbreviations: Glc-URA (glucuronic acid), Gal-URA (galacturonic acid), Man-URA (mannuronic acid). **(C)** Mole% of glucosamine in total moles of sugars exuded by giant kelp between developmental stages and seasons. Green and gold colors represent samples from mature and senescent kelp, respectively and are consistent for all three panels. Boxplots: The top and bottom border of each box represent the 25^th^ and 75^th^ percentiles, the horizontal line inside each box represents the median and the whiskers represent the range of the points (1.5*IQR) excluding outliers. Outliers are shown by black points outside the whiskers. Colored points show sample values for each boxplot.

### Seasonal and developmental stage-driven changes in exudate carbohydrate composition

Hydrolyzable carbohydrates averaged 10.3 ± 4.9% of giant kelp exudate carbon across all incubations, regardless of season or age. We observed a significant difference in the carbohydrate sugar content of kelp exudates between the mature and senescent developmental stages (Figure 2B; PERMANOVA, R^2^ = 0.14, p < 0.001) and between seasons (PERMANOVA, R^2^ = 0.07, p = 0.006). Differences in sugar content between developmental stages were driven by the change in mole% of fucose and mannuronic acid (Figure 2B) which made up, on average, 47% and 5% of the sugars exuded in the mature phase, respectively and 32% and 33% of the sugars exuded in the senescent phase, respectively (Supplemental Figure 5). Man-URA had the largest change in mole% of all sugars between the mature (5%) and senescent (34%) phases. In mature kelp blade exudates, the molar percentage of glucosamine increased significantly from summer (3%) to spring (13%), demonstrating the largest seasonal change (Figure 2C; t.test, p < 0.001).

### Microbe-Carbohydrate dynamics

Coincident with altered exudate carbohydrate composition, we observed significant differences in the giant kelp microbiome between seasons and developmental stages. In both seasons we observed a significant increase in bacterial diversity with age (Ordinary Least Squares regression; Spring: R^2^ = 0.82, p < 0.001; Summer: R^2^ = 0.23, p = 0.01) although the spring cohort had lower initial diversity (Figure 3A). Multivariate analysis demonstrated that the composition of giant kelp microbiome was significantly different between season (PERMANOVA, R^2^ = 0.26, p < 0.001) and developmental stage (PERMANOVA, R^2^ = 0.18, p < 0.001) and there was a significant interaction between season and developmental stage (Figure 3B; Two-way PERMANOVA, R^2^ = 0.10, p < 0.001). Family level shifts between mature and senescent kelp were largely driven by a significant increase in *Flavobacteriacea*, and *Arenicellaceae*, *and Rhodobacteraceae* and a significant decrease in Granulosicoccaceae, *Pirellulaceae, and Hyphomonadaceae* (Figure 4, ANOVA, FDR-adjusted p-value < 0.05, Supplemental Table 1). Family level changes were less pronounced between seasons (Supplemental Figure 6, Supplemental Table 2). At the ASV level, we identified 371 unique taxa that varied significantly in relative abundance between season or developmental stage, most of which were from the phyla Actinobacteriota, Bacteroidota, Planctomycetota, Proteobacteria (Gammaproteobacteria in particular) and Verrucomicrobiota (FDR-adjusted p-values <0.01, DEseq2, Supplemental Data 2, 3). Of these 371 ASVs, 44 had a relative abundance > 1% in more than 10% of samples. We consider these 44 ASVs to be abundant and common microbial taxa in the giant kelp microbiome. Hierarchical clustering was used to identify relationships between these 44 abundant ASVs and the mole% of sugars exuded by giant kelp based on Spearman’s rank correlation coefficients.

**Figure 3.**
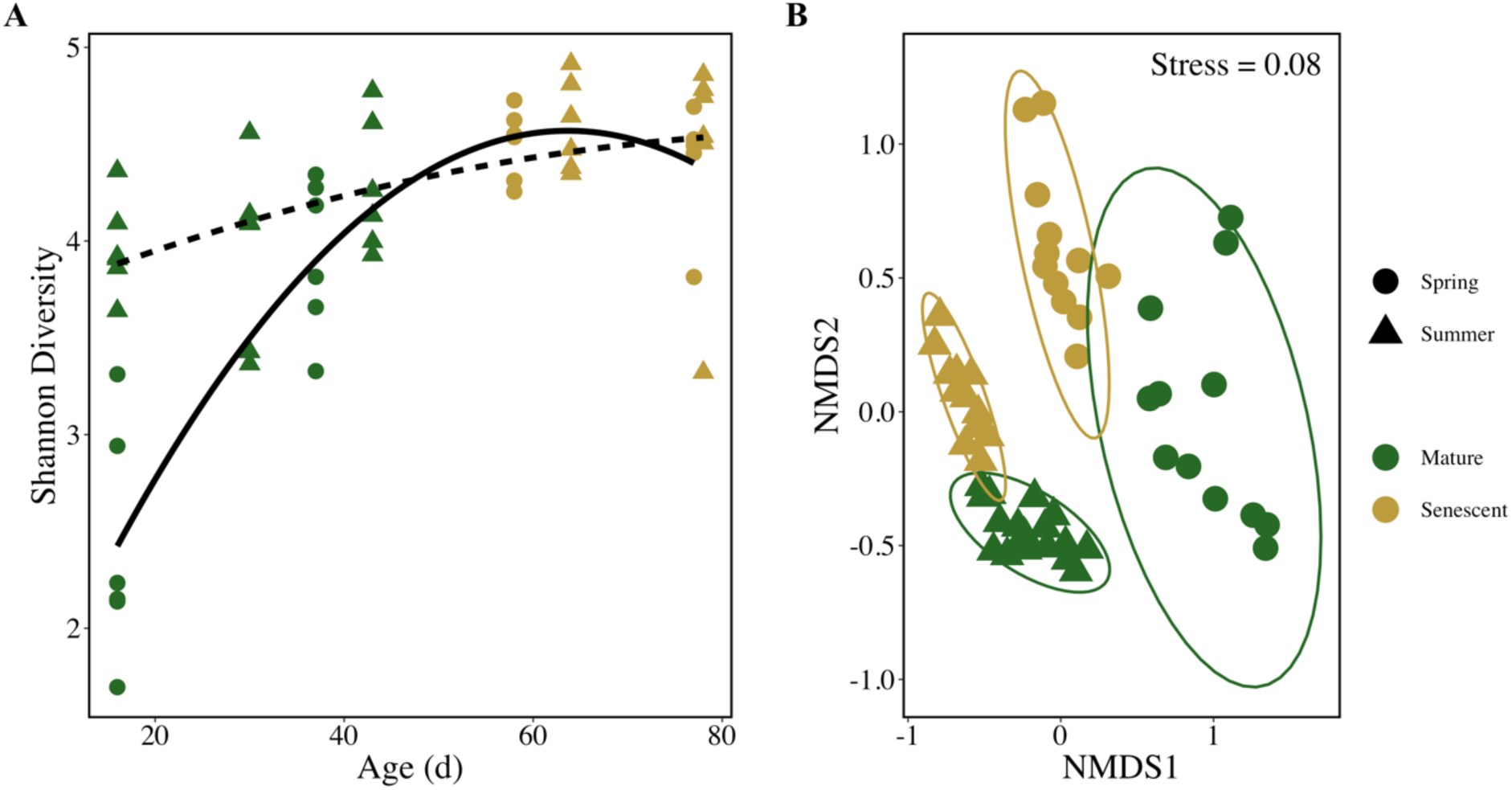
Giant kelp microbiome alpha and beta diversity between seasons and development stage. **(A)** Shannon diversity over age of the blade between spring (circles) and summer (triangles). Solid and dashed lines show the significant 2^nd^ order polynomial relationship between the giant kelp microbiome Shannon diversity and age in the spring and summer cohorts, respectively. **(B)** NMDS plot showing significant difference in the microbial community composition between seasons and development stage. NMDS ordinations used Bray-Curtis dissimilarities from arcsine-transformed 16S rRNA amplicon relative abundances. The legend applies to both panels.

**Figure 4.**
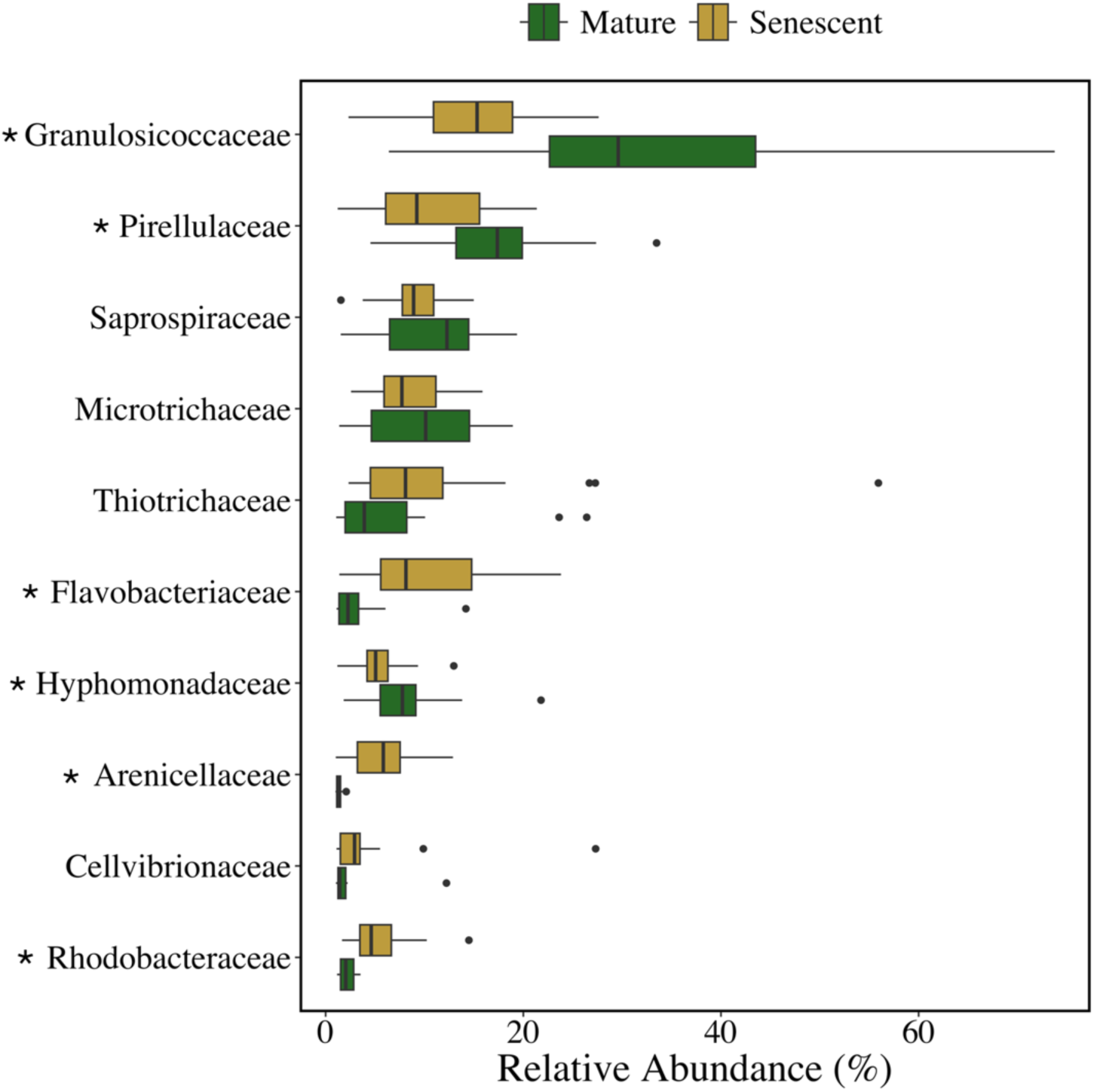
Relative abundance of bacterial clades at the family level on the surface of giant kelp blades between mature and senescent developmental stages. Shown are the top 10 families with a collective relative abundance > 1% during either developmental stage. The left and right border of each box represent the 25^th^ and 75^th^ percentiles, the vertical line inside each box represents the median and the whiskers represent 1.5*IQR. Outliers are shown by points beyond the whiskers. Asterisks next to family names indicate significant differences in the relative abundances between developmental stages (ANOVA, FDR-adjusted p-value < 0.05).

For 17 of the 44 ASVs there was at least one significant correlation (Spearman’s rank correlation, *ρ* > 0.5, FDR-adjusted p < 0.05, Supplemental Table 3 & 4) with the mole% of a sugar in giant kelp exudates. These ASVs grouped into two clusters, primarily based on their correlation with fucose, glucosamine, and mannuronic acid (Figure 5A). The first cluster (Cluster A) included ASVs whose relative abundance was positively correlated with the release of carbohydrates rich in fucose and/or glucosamine or negatively correlated with mannuronic acid (Supplemental Table 3). Members of the family *Pirellulaceae* from the phylum Planctomycetota accounted for 3 of the 6 ASVs in Cluster A. Cluster A also contained two ASVs from the family *Granulosicoccaceae* in the phylum Proteobacteria (Figure 5B).

**Figure 5.**
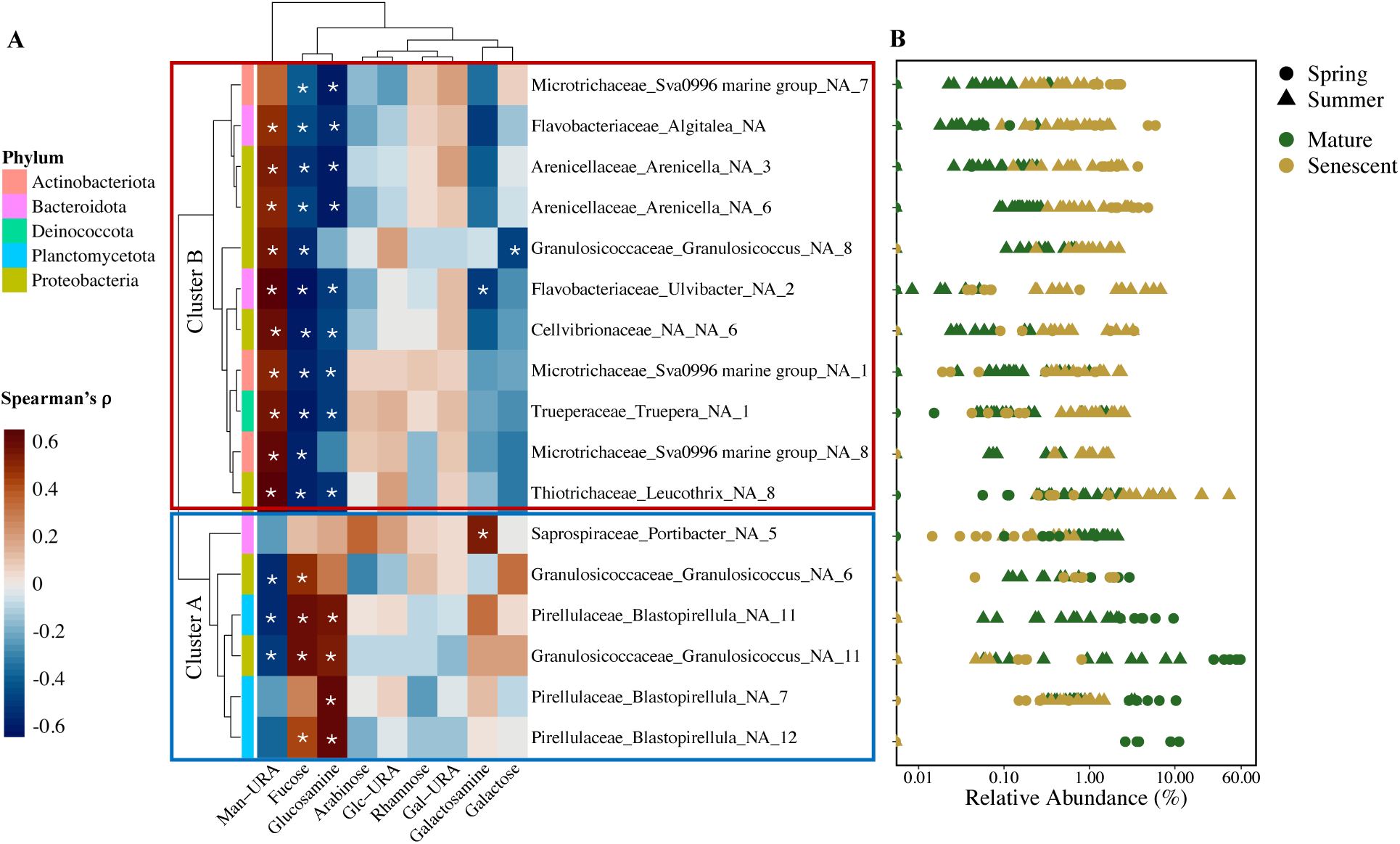
Abundant microbes are differentially correlated to the sugar content of giant kelp exudates. **(A)** Heatmap showing the Spearman’s rank correlation coefficient between the relative abundance of differentially abundant ASVs (n = 17) with at least one significant positive or negative correlation (Spearman’s rank correlation, *ρ* > 0.5, FDR-adjusted p-value < 0.05) with the mole% of a sugar monomer in giant kelp exudates. The strength and direction of the Spearman’s ranked correlation are shown by the color ramp. Significant correlations are show by the white asterisk. Hierarchical clustering shows two main clusters of ASV-sugar correlations. Cluster A is shown in the blue rectangle, and Cluster B is shown in the red rectangle. The phylum of each ASV is shown by the row color at the tip of each branch. Abbreviations: Glc-URA (glucuronic acid), Gal-URA (galacturonic acid), Man-URA (mannuronic acid). **(B)** Range in the relative abundance of each differentially abundant taxa in all kelp microbiome samples across season and physiological state. Season and physiological state of the kelp from which the sample was taken is shown by the point color and shape.

The second cluster (Cluster B) was characterized by a significant positive relationship between ASVs and the release of carbohydrates rich in mannuronic acid and/or a negative correlation with the release of carbohydrates rich in fucose and/or glucosamine (Supplemental Table 4). This cluster included members of the families *Flavobacteriaceae, Cellvibrionaceae, Arenicellaceae, Trueperaceae, Thiotrichaceae and Granulosicoccaceae*.

### Planctomycetota growth on model carbohydrates

We cultivated a bacterial isolate from the surface of a giant kelp blade belonging to the phylum Planctomycetota (family *Pirellulaceae,* genus *Rhodopirellula*) (Figure 6A & 6B) that we used in subsequent growth assays on model carbohydrates. We confirmed the ability of *this Rhodopirellula sp*. to grow on different carbohydrates, including those from kelp (Figure 6C). We observed no growth in our sterile media controls, or the carbohydrate amended media without cells. In treatments with the the isolate added we observed no cell growth in unamended treatments. The isolate grew on both *N*-acetyl glucosamine and fucoidan, but not alginate.

**Figure 6.**
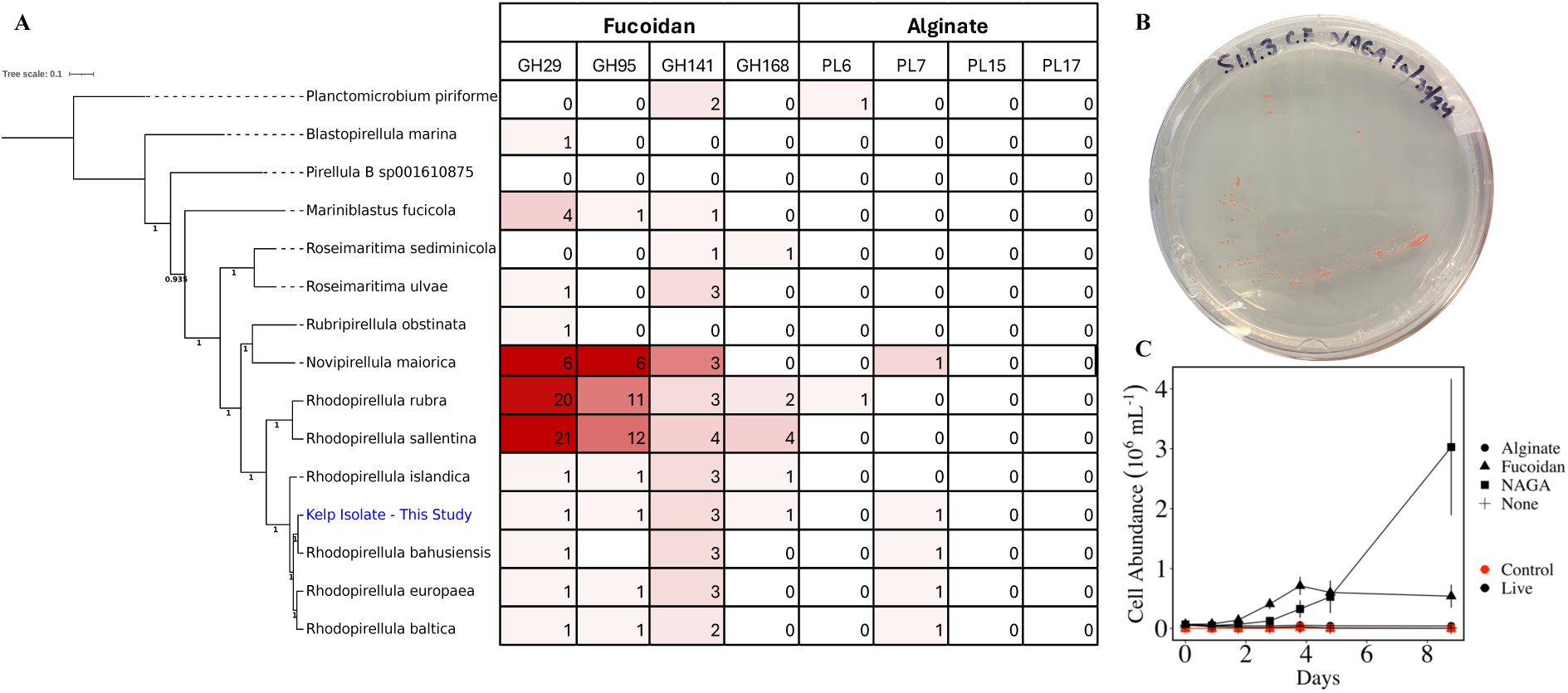
The Planctomycetota family *Pirellulaceae* are fucoidan, not alginate consumers. **(A)** Phylogenetic tree of Planctomycetota isolates within the *Pirellulaceae* family. The isolated strain from this study is highlighted in blue text. The tree was rooted with one sequence from the Planctomycetota order Planctomycetales (family *Planctomycetaceae*), located at the top of the tree. Bootstrap scores are shown at the nodes. Paired with the phylogenetic position of each isolate are the number of glycoside hydrolases (GH29, GH95, GH141, GH168) and polysaccharide lyases (PL6, PL7, PL15, PL17) involved in fucoidan and alginate degradation, respectively, predicted in their genomes. **(B)** Isolation of *Rhodopirellula* sp. on an agar plate. **(C)** Isolate growth on model carbohydrates. Control incubations (red symbols) are sterile incubation media with all carbohydrate amendments (shapes). NAGA is *N-*acetyl glucosamine. Live incubations (black symbols) show the isolate’s growth capability in response to amended carbohydrates. Points and error bars are the mean ±1 standard deviation cell abundance from triplicate incubations.

## Discussion

### Nutrient availability and age shape kelp physiology

Giant kelp physiology between and within cohorts responded to seasonal nitrogen availability and age-related senescence. At our study site, giant kelp growth is primarily limited by inorganic nitrogen availability, which is depleted in the summer, but is replenished in the spring due to coastal upwelling (Figure 1) [71, 72]. Giant kelp adjusts to periods of low nitrogen availability by adjusting its tissue nitrogen content, resulting in elevated tissue C:N ratios [71] . In our study, the average percent tissue-N of mature giant kelp in the summer (0.91 ± 0.06%) was below the minimum value (0.98%) recorded in the SBC-LTER time-series of kelp at our study site, whereas the percent tissue-N in the spring was, on average (2.6 ± 0.14%), equal to the average for our study site (2.7 ± 0.8%) [72] .In this context, our summer sampling captured kelp in a period of nitrogen depletion (Tissue-N < 1%) which is suggested to result in diminished growth rates and kelp survival [32].

In addition to seasonal fluctuations in physiology, giant kelp undergoes progressive senescence, a rapid decline in developmental stage in response to age [73]. Our results align with previous studies indicating that senescence occurs independently of environmental conditions and begins when kelp tissue age exceeds approximately 50 days [29]. These age-related physiological changes were associated with a decrease in tissue-N (Figure 2A) and changes in the composition of dissolved carbohydrates released from giant kelp (Figure 2B & 2C).

### Giant kelp microbiome composition

The relative abundance of major epibiont bacterial families at different stages of kelp development is broadly consistent with observations of kelp microbiomes over tissue age and between healthy and stressed or senescent tissue (Figure 4) [35, 41, 42, 74, 75]. Likewise, the increase in diversity of the microbiome with age is consistent with previous observations [74]; however, to our knowledge this is the first report of significantly lower Shannon diversity in young kelp in the spring compared to the summer (Figure 3A). It has been proposed that the kelp microbiome is established through environmental acquisition over vertical transmission [76]. Therefore, exudates likely play a major role in the establishment and transformation of the kelp microbiome. Kelps continuously shed and replace tissue year-round [77] and factors that affect exudate composition, such as nitrogen availability, may have consequences for the biofilm composition of new tissue, as has been proposed for developing terrestrial plant rhizospheres [7].

### Exudate and microbiome composition dynamics

Simultaneous changes in the carbohydrate composition of kelp exudates and microbial taxa suggest an important role in exudate composition in shaping the kelp microbiome. These associations further revealed that extrinsic factors, such as nitrogen availability, and intrinsic factors such as development stage regulate the composition of the giant kelp exudates. Notably, we observed an enrichment of glucosamine, a nitrogen-containing sugar, in carbohydrates exuded by mature kelp in the spring compared to the nitrogen-depleted kelp in the summer (Figure 1C). A similar pattern has been observed in the root exudates of plants, whereby the addition of inorganic nitrogen results in the enhanced release of nitrogen containing metabolites relative to roots grown under nitrogen-limiting conditions. Baker et al., (2024) [7] found that the release of nitrogen-rich metabolites by plant roots selected for a less diverse and unique microbial community compared to plants grown in nitrogen-limited conditions, a pattern we also observed in the kelp microbiome (Figure 2). Glucosamine is a common sugar in the mucus produced by many organisms, including corals and humans [15, 18, 78], and has been shown to serve an important role in regulating attachment and biofilm formation by bacteria [79, 80]. Because kelp exudates are constantly sloughed into the surrounding seawater, they must be replaced to maintain their function in the extracellular matrix (mucilage). Therefore, the cost of exuding nitrogen containing compounds that must be continuously replaced may be too high to be maintained under nitrogen-depleted conditions experienced by giant kelp in the summer.

Fucose was the most abundant sugar in the carbohydrates released by mature giant kelp in both seasons, consistent with previous studies of brown macroalgae [17, 51, 52]. The presence of fucose in brown macroalgal exudates is due to the secretion of fucoidan, a water-soluble, sulfated polysaccharide that is deposited onto the exterior of the thallus through secretory canals to build the kelp’s extracellular mucilage, which is continuously sloughed into the surrounding seawater [81, 82]. Fucoidan lacks nitrogen which may explain its consistent dominance in kelp exudates across a range of tissue-N content. Several studies document the potential defensive properties of fucoidans [69, 79, 83] and its proposed defensive role is supported by its observed recalcitrance to bacterial degradation [84].

In both seasons, there was an increase in the mole% of mannuronic acid as kelp blades aged and senesced (Figure 1B). Mannuronic acid is an indicator of the macroalgal polysaccharide alginate, which comprises up to half of kelp biomass [85]. Microbial taxa positively correlated to the mole% of mannuronic acid included members of the family *Flavobacteriaceae* and several Gammaproteobacteria, including one member of the family *Thiotrichaceae* (Figure 5A, Cluster B). Flavobacteria and Gammaproteobacteria are archetypal fast-growing carbohydrate degraders with adaptations to grow attached to surfaces and are enriched in alginate and oligoalginate lyases [86–89]. Using DNA stable isotope probing with ^13^C-labeled alginate, Thomas et al., (2021) [86] demonstrated the main consumers of alginate in coastal seawater were bacteria from the families *Flavobacteriaceae*, *Alteromonadaceae*, *Thiotrichaceae* clades. The relative abundance of *Flavobacteriaceae* and *Thiotrichaceae* in kelp biofilms is typically higher on stressed or senescent kelp tissue [35, 75] as observed in this study (Figure 4), and in the early stages of kelp biomass decay these groups dominate alginate hydrolysis [87]. From our observations, we hypothesize that as kelp begins to senesce, the hydrolysis of insoluble alginate polymers promotes the growth of alginate degraders whose hydrolytic enzymes further solubilize kelp tissue into DOC. This dynamic would explain the enhanced solubilization of kelp tissue with age that we observed in a complementary study [90].

The senescence-associated release of alginate is likely initiated by early alginate degrading “pioneer” bacteria whose alginate lyases release alginate oligomers into the environment, facilitating the growth of alginate “harvesters” and “scavengers” [91] Notably, these potential alginate degraders were observed at low relevant abundance in the mature kelp microbiome (Figure 5B), suggesting some process limited their growth until kelp entered its senescent phase. The mechanisms of kelp senescence are not well understood; however, senescence of giant kelp blades in both seasons occurred after 50 days of age, following linear decline in photosynthetic rates (see our complementary study [90]), suggesting intrinsic regulation of senescence [92]; although, we cannot completely rule out that associated bacteria contributed to senescence.

### Potential mechanisms behind carbohydrate-microbe associations

Without further study of their native composition, we can only speculate about the structure and function of the carbohydrates released by giant kelp. However, the role of carbohydrates secreted to the outer membrane layers of animals and phytoplankton is well established. For example, in bacterial biofilm formation, carbohydrates tethered to the cell surface of a host serve as recognition sites for carbohydrate-protein binding proteins (lectins) that act in concert with larger attachment proteins (adhesins). These lectin-adhesin complexes mediate the attachment of bacteria, including pathogens to the host surface [93]. In a model bacterium-diatom association, the lectin *Mp*PA14 exhibits a strong binding affinity for fucose and *N*-acetyl glucosamine residues [79]. Guo et al., (2021) demonstrated that *Mp*PA14 and the *Mp*PA14 lectin homolog from *Vibrio cholera* were inhibited and unable to bind to diatom cells when exposed to high concentrations of free fucose. A strong binding affinity of *Mp*PA14 was also observed for fucoidan, a fucose-rich polysaccharide released by brown macroalgae. These data indicate a potentially conserved defensive role of fucose-containing carbohydrates exuded by members of the Stramenopile lineage, such as diatoms and kelps.

Giant kelp may release high concentrations of fucose-rich carbohydrates to disrupt biofilm formation by potential pathogens. Indeed, all taxa from Cluster B (Figure 4A), which includes several taxa associated with macroalgal tissue degradation, such as Flavobacteria, exhibited a negative correlation with the fucose content of the exuded carbohydrates (Spearman’s ranked correlation, FDR-adjusted p < 0.05; Supplemental Table 4). This was surprising, as previous metagenomic-based studies predict that members of the Bacteroidetes phylum, especially *Flavobacteraceae*, are specialized to degrade fucoidans [94, 95]. Therefore, other factors, in addition to carbohydrate substrate preferences may contribute to the structuring of the kelp microbiome; possible factors could be the production of secondary metabolites by Planctomycetota [63, 96] or the anti-biofilm property of fucoidan [79]. Regarding the former, AntiSMASH analysis of our isolated *Rhodopirellula* genome annotated two terpene biosynthetic gene clusters, two type I polyketide synthases (PKS), one type III PKS, and hybrid type I PKS/terpene and type I/non ribosomal peptide synthases (Supplemental Figure 7). This is consistent with other isolates from the *Pirellulaceae* family [63] and suggests their abundances in kelp biofilms may be due to a combination of their ability to degrade fucoidan (Figure 6C) and their production of defensive secondary metabolites. Experiments to address secondary metabolite production by this isolate are currently underway.

Glucosamine may serve as an important carbon and nitrogen source for the growth of microbes beneficial to giant kelp. In the marine environment, glucosamine is more commonly found in its acetylated form (*N*-acetyl glucosamine) [97]. Our carbohydrate measurements require an acid hydrolysis step, which cleaves the acetyl group resulting in glucosamine. Therefore, our glucosamine measurements reflect the contribution of both *N*-acetyl glucosamine and glucosamine residues. The microbial taxa that positively correlated with glucosamine included members from the family *Pirellulaceae* within Planctomycetota, and the *Granulosicoccaceae* family of Proteobacteria (Cluster A, Figure 5A, Supplemental Table 3). Cultured strains of Planctomycetota such as *Pirellula* sp. strain 1 can metabolize *N*-acetyl glucosamine (NAGA) as their sole carbon and nitrogen source [98], and in a comparison of 79 cultured members of Planctomycetota, *N*-acetyl glucosamine was found to be the most efficient carbon source for their isolation [63]. Moreover, previous work suggests *N*-acetyl glucosamine may trigger biofilm formation by some members of Planctomycetota, such as *Rhodopirellua baltica* [99].

Planctomycetes are dominant members of healthy macroalgal microbiomes [35, 42, 100] and it has been proposed they alter the composition of biofilms through the production of secondary metabolites such as stieleriacines [96]. We demonstrate that a *Rhodopirellula* strain, isolated from the surface of a giant kelp blade, can grow on fucoidan and NAGA, but not alginate (Figure 6C). This carbohydrate substrate preference is consistent with our observed correlations between the relative abundance of members of the Planctomycetota family *Pirellulaceae* in the kelp microbiome and the sugar content of giant kelp exudates (Figure 5A, Cluster A). Although only one isolate was tested in this study, the ability for growth on fucoidan but not alginate is broadly consistent with the metabolic potential of other isolated members of the family *Pirellulaceae* (Figure 6A). Further, several recent studies also confirm that the isolates from the *Pirellulaceae* family degrade fucoidan [38, 101]. We also note that even though some *Pirellulaceae* genomes, including our isolate genome, contained copies of the alginate lyases PL6 or PL7, none contain the oligoalginate lyases PL15 or PL17, which degrade oligoalginates into monomers for metabolism (Figure 6A) [91]. We hypothesize that exudation of specific carbohydrates, such as fucoidan serves to enrich their microbiome in commensal or beneficial microbes such as Planctomycetota. Such carbohydrate-microbe synergy is consistent with observations of plant rhizospheres, corals and the human gut [15, 78, 102–104].

## Acknowledgments

We thank all past and current members of the Santa Barbara Coastal LTER who generated necessary background data for this study. We thank all Carlson lab members, A. Santoro and D. Siegel for their constructive comments and discussion of the data and manuscript. We thank P. Thieringer for his advice on whole genome assembly and comparative genomics. This project was funded by the Department of Energy’s Advanced Research Project Agency-Energy through award DE-AR0001559 to CAC, and the National Science Foundation’s Santa Barbara Coastal LTER through award number OCE-1831937 to CAC. CJE and MM were additionally supported by the Worster Family Summer Research Fellowship awarded to CJE.

## Data Availability

DNA sequence data are available in the National Center for Biotechnology Information Sequence Read Archive under PRJNA1256130. R scripts for analysis can be found at https://github.com/chance-english.

## Supplementals

**Supplemental Table 1:** ANOVA results for differences in relative abundance of top 10 families btween mature and developmental stages shown in Figure 4. Significant results means the observed relative abundances of the family was higher in in the mature developmental stage

| family | term | df | sumsq | meansq | statistic | p.value | adjusted_p.value |
| --- | --- | --- | --- | --- | --- | --- | --- |
| Granulosicoccaceae | phys_state | 1 | 0.38 | 0.38 | 20.47 | 3.65E-05 | 3.65E-04 |
| Thiotrichaceae | phys_state | 1 | 0.03 | 0.03 | 2.82 | 0.100 | 0.144 |
| Pirellulaceae | phys_state | 1 | 0.07 | 0.07 | 17.75 | 1.01E-04 | 5.03E-04 |
| Cellvibrionaceae | phys_state | 1 | 0.00 | 0.00 | 0.23 | 0.638 | 0.638 |
| Flavobacteriaceae | phys_state | 1 | 0.04 | 0.04 | 17.37 | 1.71E-04 | 5.71E-04 |
| Hyphomonadaceae | phys_state | 1 | 0.01 | 0.01 | 7.99 | 0.007 | 0.011 |
| Saprospiraceae | phys_state | 1 | 0.00 | 0.00 | 1.21 | 0.277 | 0.344 |
| Microtrichaceae | phys_state | 1 | 0.00 | 0.00 | 1.06 | 0.309 | 0.344 |
| Rhodobacteraceae | phys_state | 1 | 0.01 | 0.01 | 16.11 | 2.71E-04 | 6.78E-04 |
| Arenicellaceae | phys_state | 1 | 0.01 | 0.01 | 15.72 | 4.03E-04 | 8.06E-04 |

**Supplemental Table 2:** ANOVA results for differences in relative abundance of top 10 families between spring and summer cohorts shown in Supplemental Figure 6.Significant results means the observed relative abundances of the family was higher in the spring.

| family | term | df | sumsq | meansq | statistic | p.value | adjusted_p.value |
| --- | --- | --- | --- | --- | --- | --- | --- |
| Granulosicoccaceae | season | 1 | 0.045 | 0.045 | 1.79 | 0.1870 | 0.255150 |
| Thiotrichaceae | season | 1 | 0.009 | 0.009 | 0.90 | 0.3487 | 0.387435 |
| Pirellulaceae | season | 1 | 0.009 | 0.009 | 1.92 | 0.1721 | 0.255150 |
| Cellvibrionaceae | season | 1 | 0.006 | 0.006 | 1.73 | 0.2041 | 0.255150 |
| Flavobacteriaceae | season | 1 | 0.036 | 0.036 | 14.10 | 0.0006 | 0.001930 |
| Hyphomonadaceae | season | 1 | 0.000 | 0.000 | 0.08 | 0.7820 | 0.781972 |
| Saprospiraceae | season | 1 | 0.056 | 0.056 | 52.15 | 0.0000 | 0.000000 |
| Microtrichaceae | season | 1 | 0.037 | 0.037 | 20.60 | 0.0000 | 0.000203 |
| Rhodobacteraceae | season | 1 | 0.007 | 0.007 | 10.69 | 0.0023 | 0.005728 |
| Arenicellaceae | season | 1 | 0.002 | 0.002 | 1.71 | 0.2005 | 0.255150 |

**Supplemental Table 3.**
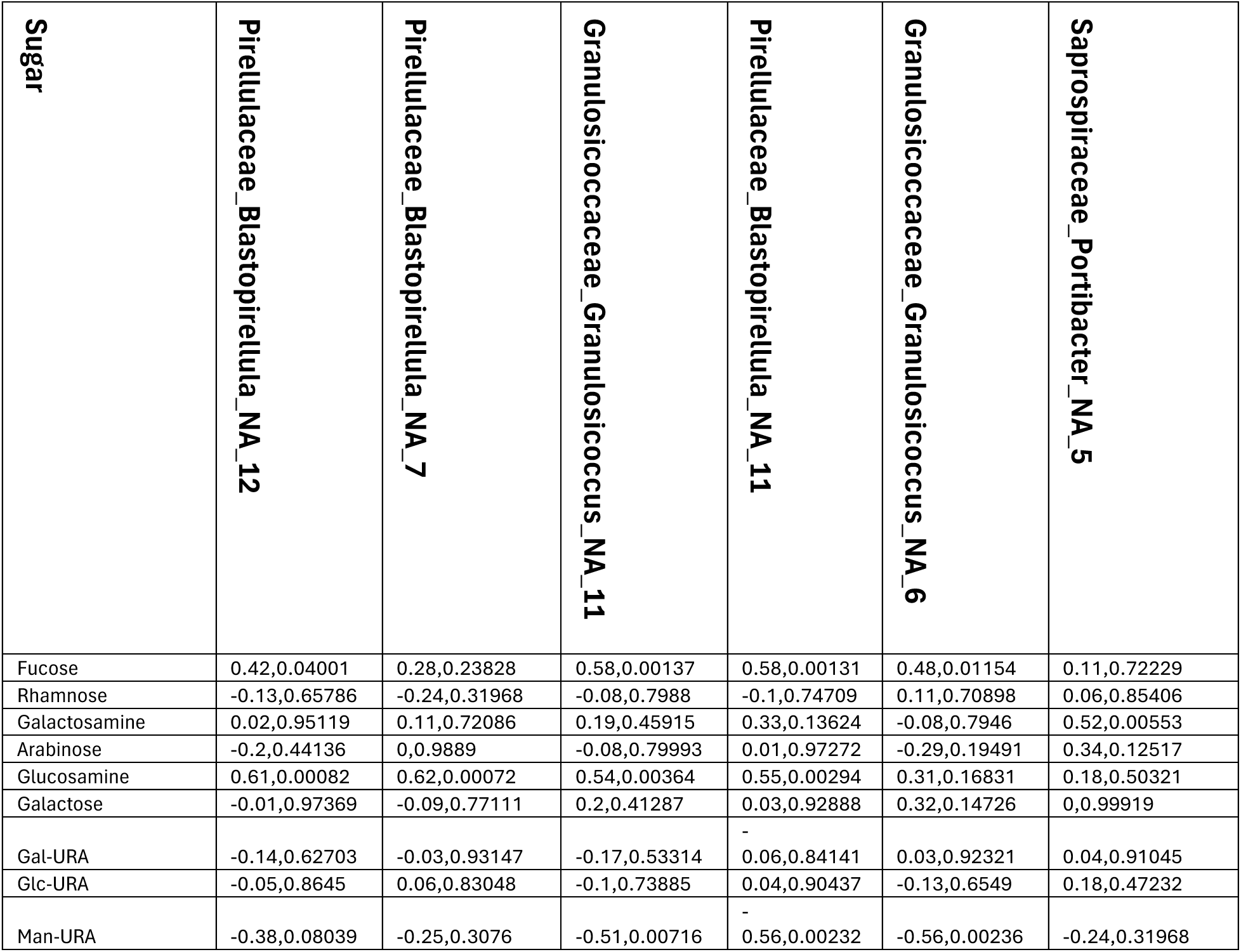
Spearman’s rank correlation rho and FDR-adjusted p-values for abundant ASVs and sugar mole% in Cluster A (. **Figure 5A).**

**Supplemental Table 4.**
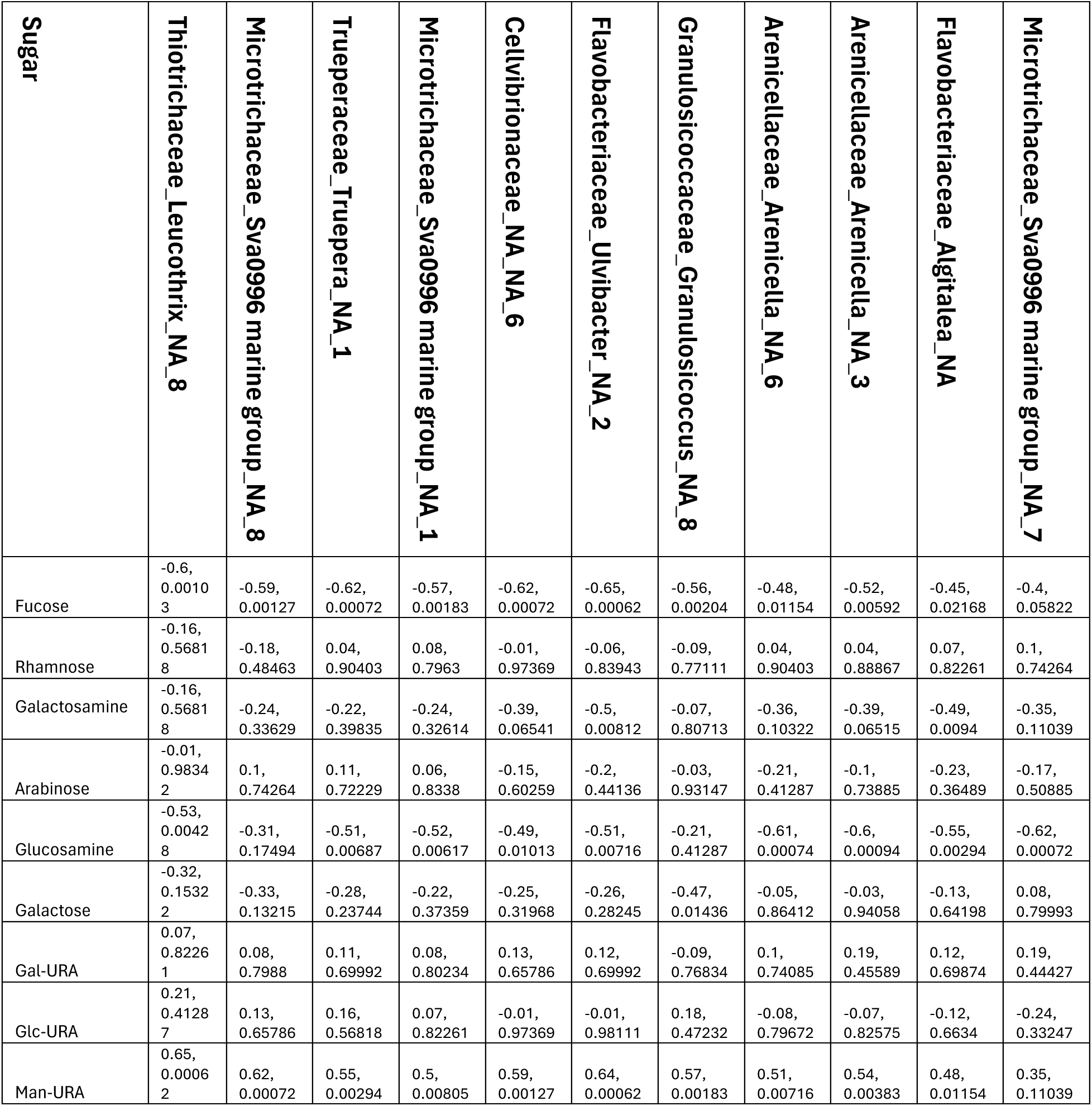
Spearman’s rank correlation rho and FDR-adjusted p-value for abundant ASVs and sugar mole% in Cluster B (. **Figure 5A).**

**Supplemental Figure 1.**
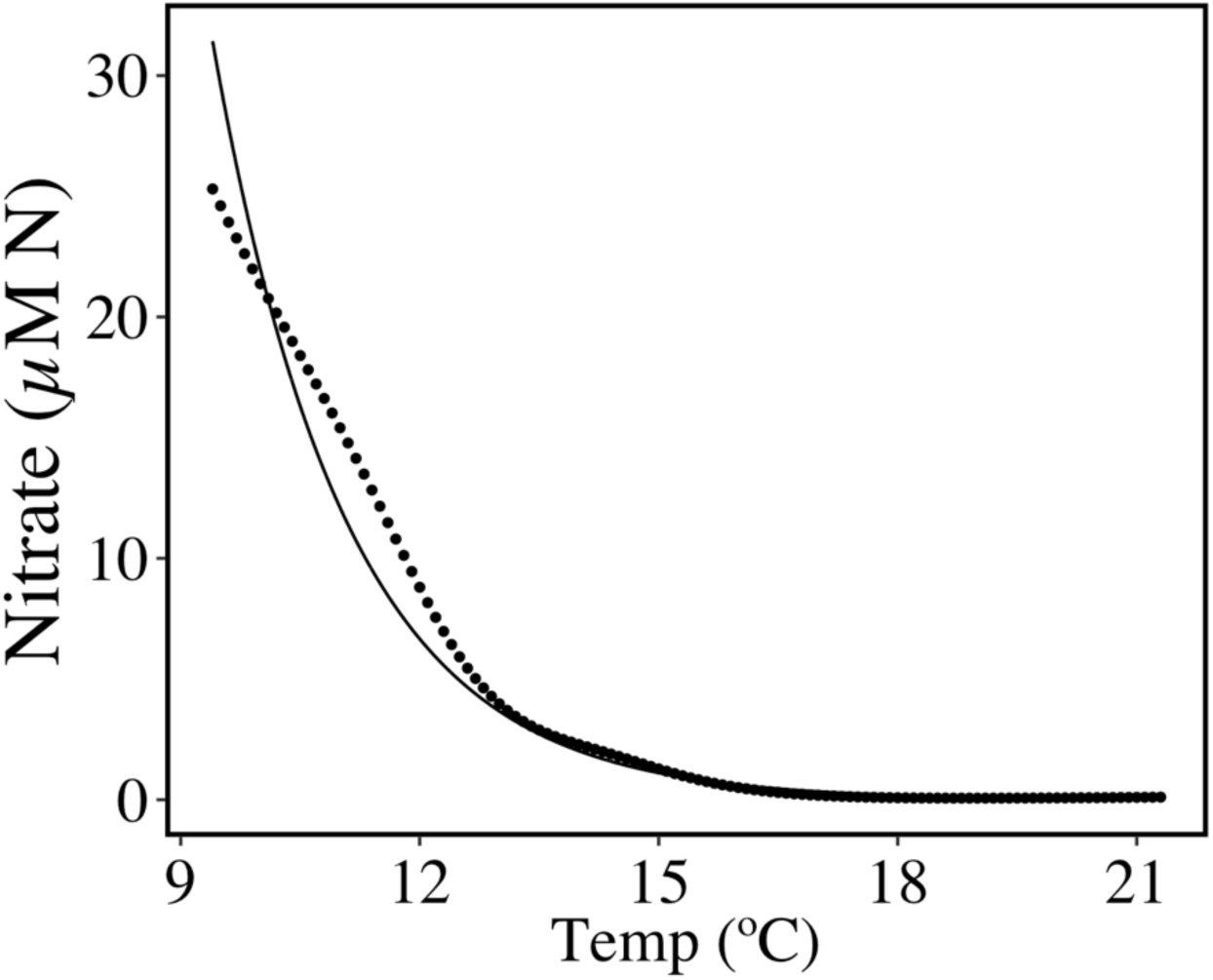
Temperature to nitrate relationship for inshore water in the Santa Barbara Channel. Points are from the T2N look up table compiled by Snyder et al. 2020 . Solid line is the exponential regression used to calculate nitrate at Mohawk Reef at our study site: Nitrate (µM N) = 8565.5 * e^(-0.597*Temp(°C))^, R^2^ = 0.95.

**Supplemental Figure 2.**
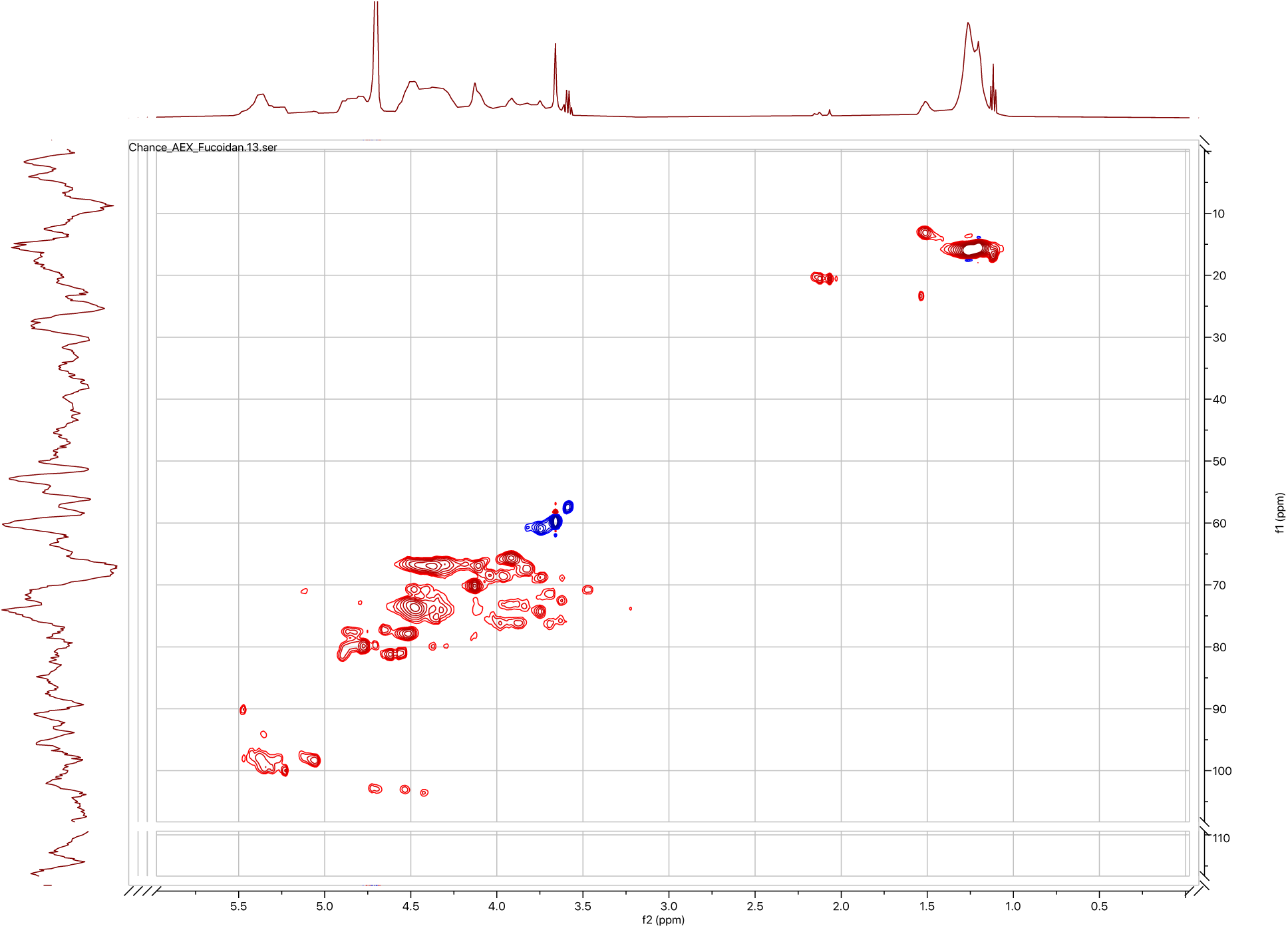
1H-^13^C HSQC NMR spectrum of IEX purified fucoidan from *Macrocystis pyrifera*. Correlations showing carbons with an even number of protons are blue and those with an odd number of protons are shown in red.

**Supplemental Figure 3.**
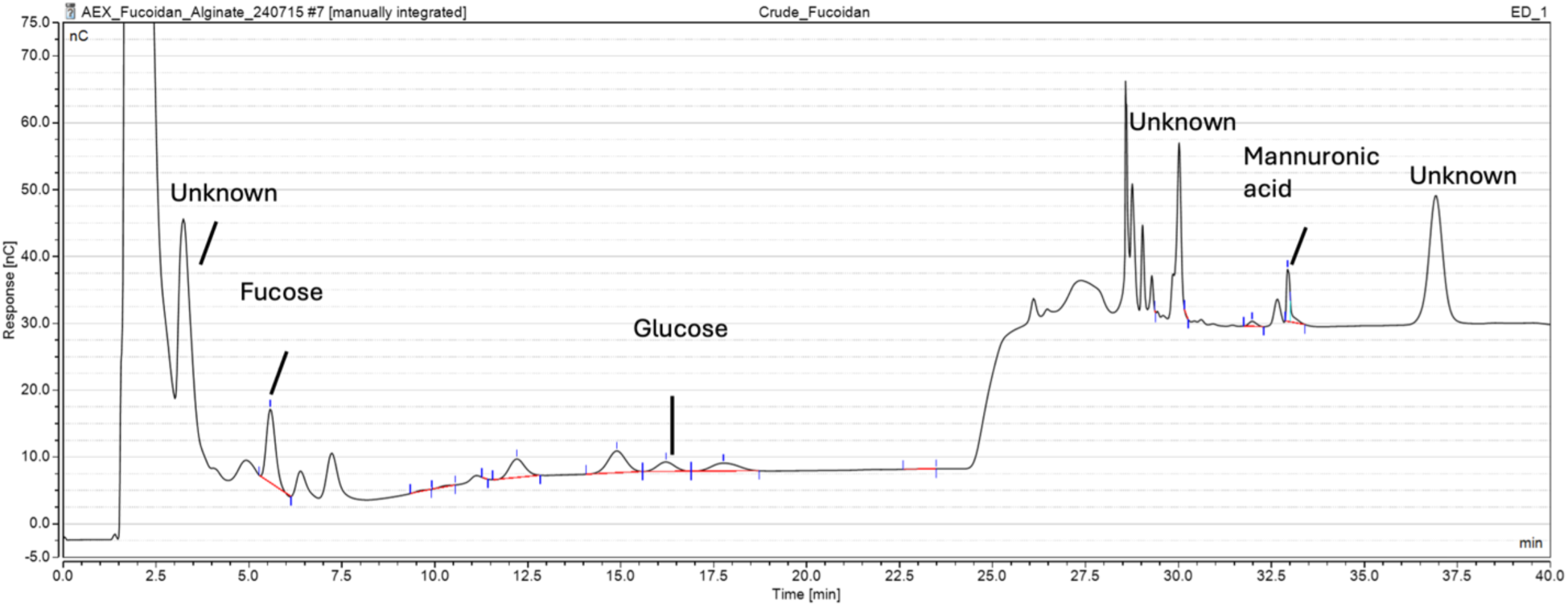
HPAEC-PAD chromatogram of crude fucoidan from supplier. Note that peak area for fucose is relatively equal to areas from sugars not found in fucoidan (glucose, mannuronic acid) compared to fucose in IEX-purified fucoidan (Supplemental Figure 4). Also, there are large contributions from unknown peaks, possibly contaminating sugars in the sugar alcohol range (0 - 5 minutes) and acidic sugar range (30 - 40 minutes).

**Supplemental Figure 4.**
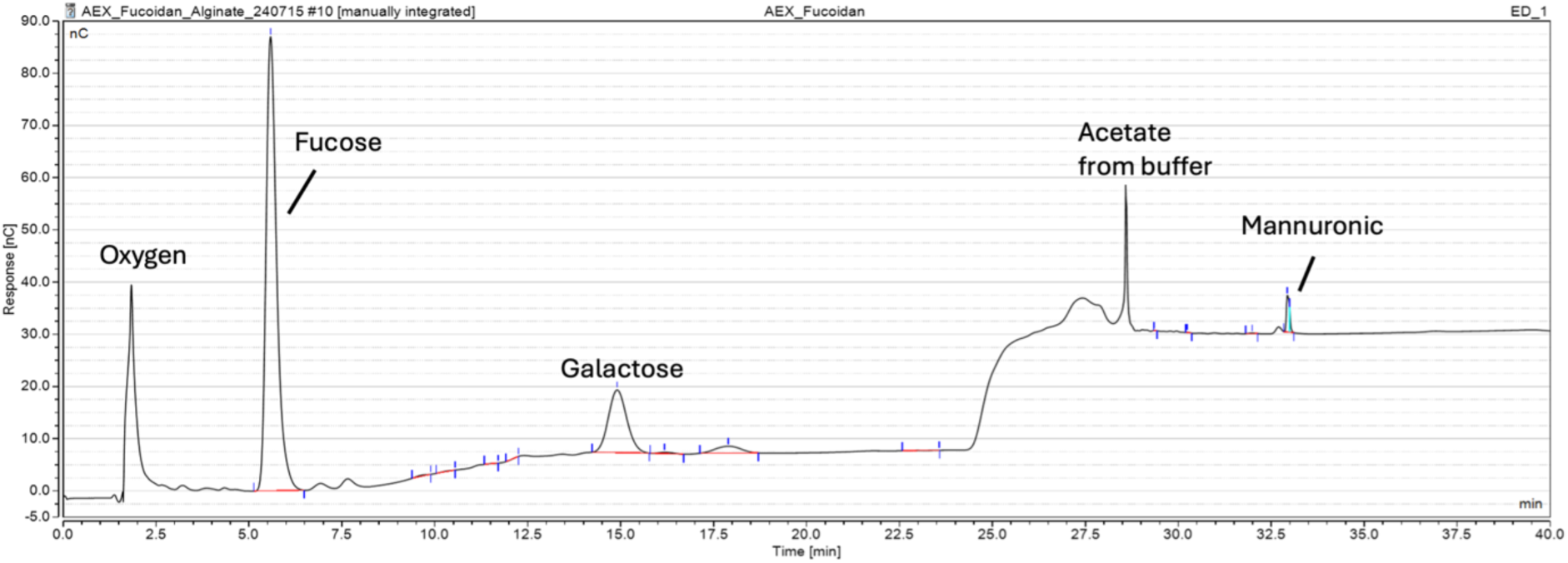
HPAEC-PAD chromatogram of IEX purified fucoidan. Note that peak area of fucose enhanced compared to small remaining contaminating sugars (mannuronic acid) compared to Supplemental Figure 3. Contribution of galactose indicates fucoidan from *M. pyrifera* is likely a heterofucan. The two prominent peaks at the beginning and middle of the chromatogram are from dissolved oxygen and acetate from the sample and elution buffer, respectively.

**Supplemental Figure 5:**
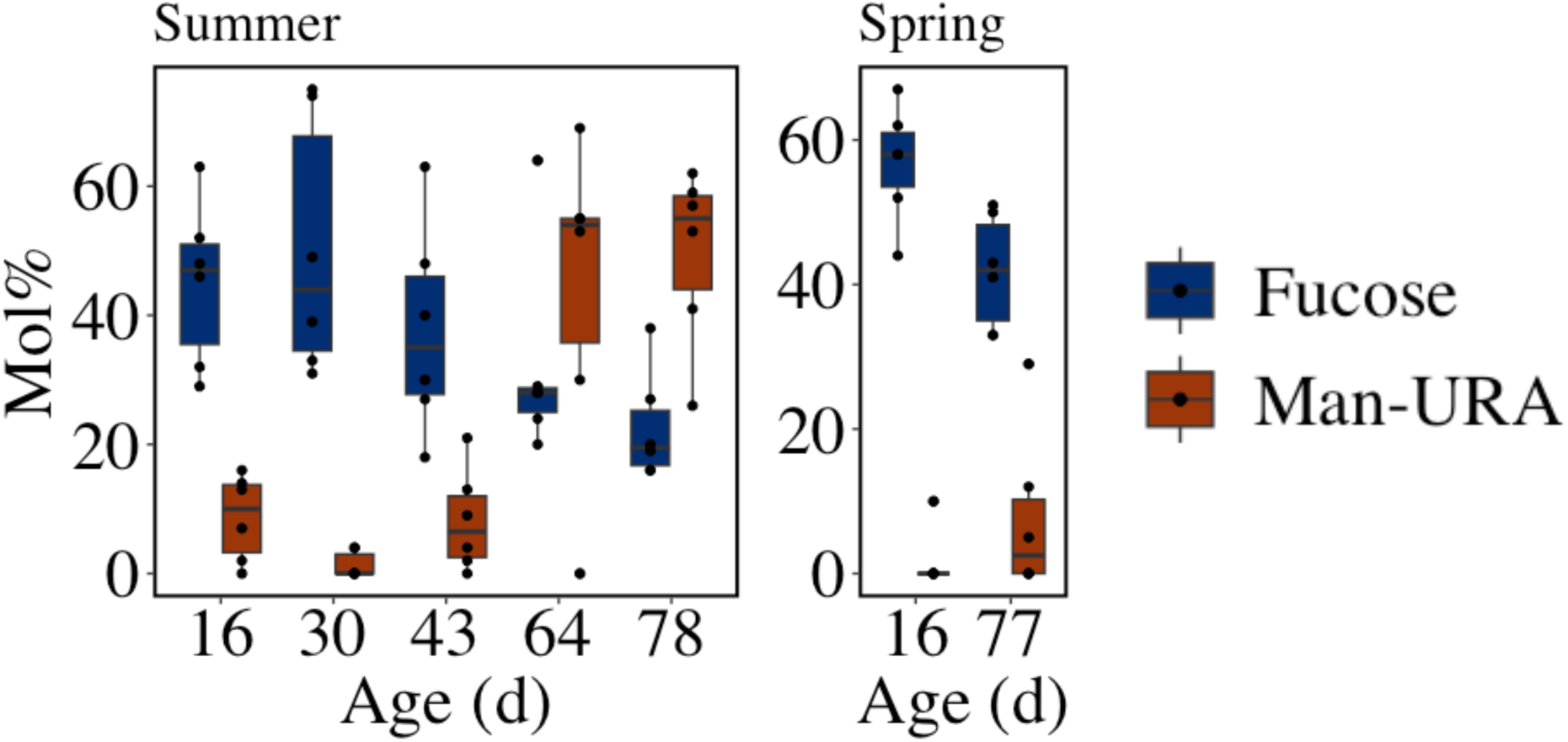
Mole% of fucose and mannuronic acid (Man-URA) across age and and seasons. **(A) Summer (B) Spring.** The top and bottom border of each box represent the 25^th^ and 75^th^ percentiles, the horizontal line inside each box represents the median and the whiskers represent 1.5*IQR. Outliers are shown by points beyond the whiskers.

**Supplemental Figure 6:**
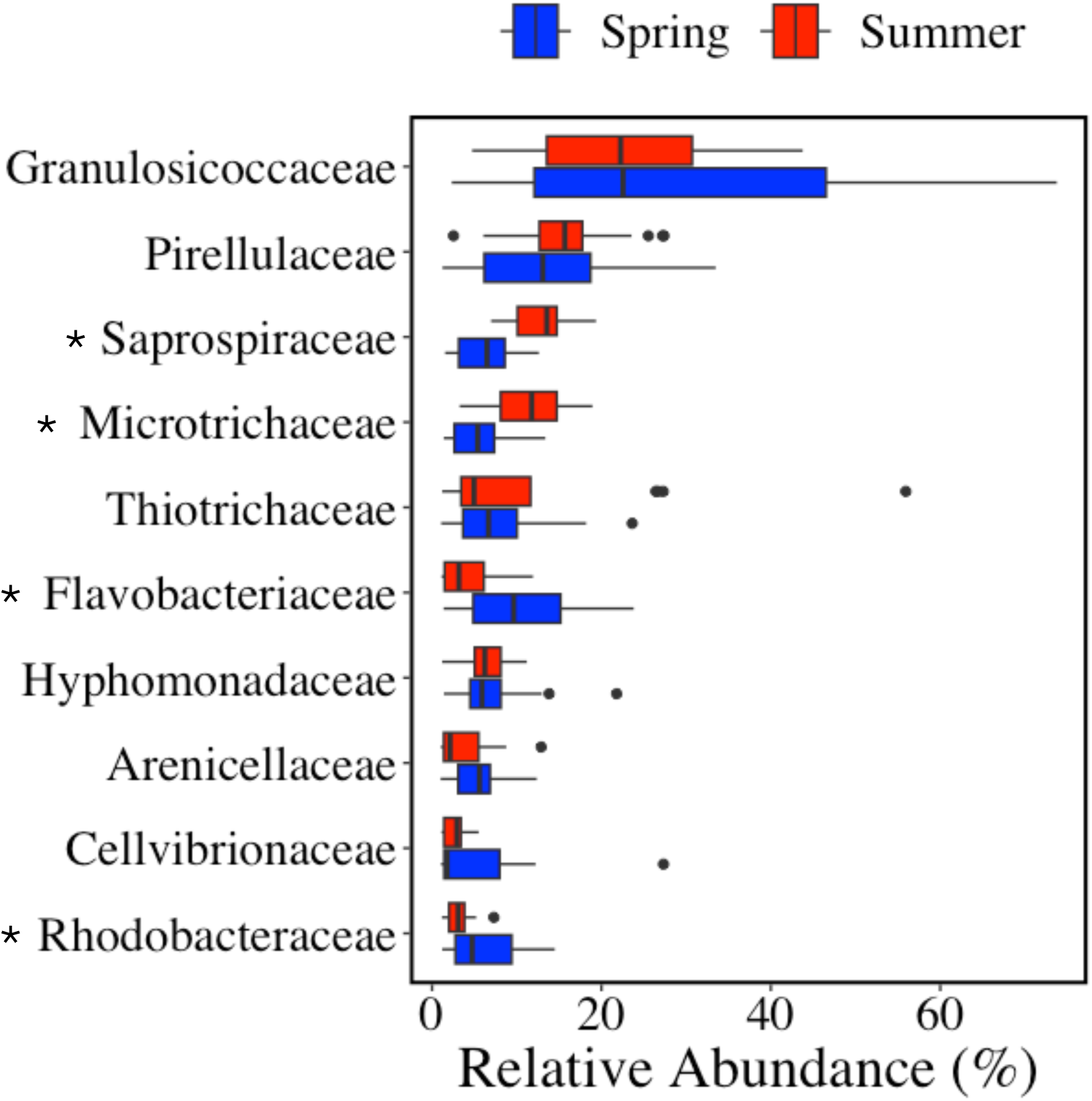
Relative abundance of bacterial clades at the family level on the surface of giant kelp blades between the spring and summer periods. Shown are the top 10 families with a collective relative abundance > 1% during either season. The left and right border of each box represent the 25^th^ and 75^th^ percentiles, the vertical line inside each box represents the median and the whiskers represent 1.5*IQR. Outliers are shown by points beyond the whiskers. Asterisks next to family names indicate significant differences in the relative abundances between seasons (ANOVA, FDR-adjusted p-value < 0.05).

**Supplemental Figure 7:**
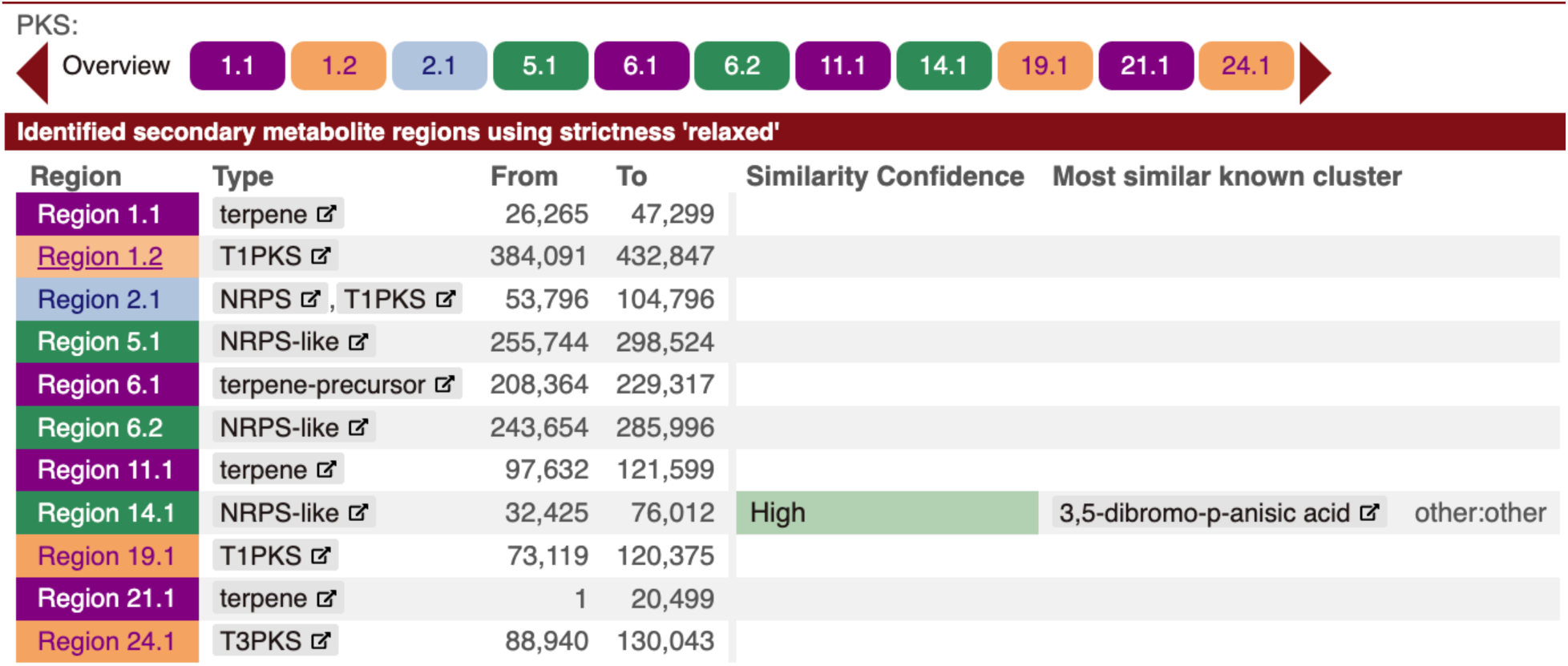
AntiSMASH analysis results of the *Rhodopirellula sp* isolate genome.

